# Shared and disease-specific human brain vascular signatures in Alzheimer’s disease, frontotemporal dementia, and Huntington’s disease

**DOI:** 10.64898/2026.07.01.735865

**Authors:** Patrycja M. Forster, Oliver Bracko, Ruslan Rust

**Affiliations:** Laboratory for Intestinal Neuroimmune Interactions, Department of Chronic Diseases, Metabolism and Ageing, Translational Research Center for Gastrointestinal Disorders, KU Leuven, Leuven, Belgium; Department of Physiology and Neuroscience, University of Southern California, 90033, Los Angeles, USA; USC Epstein Family Alzheimer’s Therapeutic Research Institute, 9860, San Diego; Zilkha Neurogenetic Institute, Keck School of Medicine, University of Southern California, 90033, Los Angeles, USA

**Keywords:** Single-cell RNA sequencing, blood-brain barrier, brain vasculature, pericytes, cell-cell communication, Alzheimer’s, dementia, Huntington’s, human

## Abstract

The blood-brain barrier (BBB) plays a central role in brain function and is increasingly implicated in neurodegenerative disease. Major neurodegenerative disorders, including Alzheimer’s disease (AD), frontotemporal dementia (FTD), and Huntington’s disease (HD), share overlapping pathological features. Yet, the extent to which these diseases converge or diverge at the level of BBB-associated cell types remains poorly understood. Here, we performed a comparative analysis of vessel-enriched human brain transcriptomic datasets across AD, FTD, and HD to define shared and disease-specific neurovascular alterations. We identify a partially conserved transcriptional signature of vascular dysfunction across all three diseases, alongside disease-specific changes in endothelial, pericyte, and perivascular cell populations. Endothelial remodeling was most prominent in capillary and venous segments, highlighting segment-specific vulnerability along the arteriovenous axis. Notably, we identified two molecularly distinct human pericyte subtypes across all three datasets and found a consistent reduction in the matrix-type pericytes (M-peri) fraction, suggesting a selective decline. Cell-cell communication analysis further revealed reorganized endothelial-pericyte signaling networks, with prominent alterations in extracellular matrix-associated pathways, including LAMININ, COLLAGEN, FN1, and NCAM, together with changes in contact-dependent and vascular signaling pathways such as NOTCH and VEGF. Together, our findings define shared and disease-specific neurovascular mechanisms across major neurodegenerative disorders and highlight BBB-associated pathways as central features of neurodegeneration, providing a framework for future diagnostic and therapeutic strategies.

## Introduction

The blood-brain barrier (BBB) is a specialized vascular interface formed by endothelial cells and supported by pericytes, smooth muscle cells, astrocytic endfeet, fibroblasts, and other immune-associated perivascular cell types. Together, they regulate the entry of toxins, immune cells, and pathogens into the brain, enable controlled transport of nutrients, metabolic substrates, and signaling molecules, and facilitate the clearance of metabolic waste from the brain into the blood^1–3^. Neuronal function depends strongly on this tightly regulated vascular environment, and disruption of BBB integrity and function has increasingly been implicated in neurodegenerative diseases^1–6^. Emerging evidence suggests that BBB dysfunction is not solely a downstream consequence of neurodegeneration but may actively contribute to neuronal, synaptic, and cognitive impairment, altered immune surveillance, and broader disease progression^7,8^.

Alzheimer’s disease (AD), frontotemporal dementia (FTD), and Huntington’s disease (HD) are among the most common neurodegenerative diseases worldwide^9^. Although these disorders are traditionally defined by distinct pathological hallmarks, including amyloid-β and tau accumulation in AD^10^, mutant huntingtin protein in HD, and tau, TDP-43, or FUS pathology in FTD, they also show substantial clinical, genetic, and neuropathological overlap. For example, TDP-43 pathology is frequently observed in AD, and inflammatory, metabolic, synaptic, and vascular changes are increasingly recognized across multiple neurodegenerative conditions^13^. Such overlap suggests that, despite disease-specific initiating mechanisms, shared tissue-level responses may contribute to disease progression.

To date, efforts to identify shared and disease-specific processes across neurodegenerative diseases have focused on either blood-derived samples^14^ or all major brain cell types^15–19^, whereas human brain vasculature cell types remain understudied. Recent single-cell and single-nucleus studies have begun to reveal the cellular complexity of the human brain vasculature in health and disease^20,21^. In AD, vascular profiling has identified disease-associated changes in endothelial and perivascular populations, including altered vascular risk-gene expression and selective vulnerability of extracellular matrix-maintaining pericytes (M-peri)^21^. In FTD linked to GRN mutations, neurovascular dysfunction has been associated with BBB-related endothelial signatures, altered pericyte coverage, fibroblast and immune activation, and increased vascular remodeling^22^. In HD, cerebrovascular single-cell analyses have revealed vascular and glial immune activation along with reduced expression of proteins important for BBB integrity^23^. Together, these studies suggest that neurodegenerative diseases affect the neurovascular unit, but they have largely been analyzed independently.

Whether neurodegenerative diseases converge or diverge at the level of brain vascular and perivascular cell types remains largely unknown. In particular, it is unclear which vascular alterations, vascular endothelial zonation changes, pericyte states, and ligand-receptor communication processes might be involved in disease progression and could be shared across disorders, and which of these reflect disease specific pathology. This question is especially relevant for pericytes, which occupy a central position within the neurovascular unit. Pericytes regulate BBB integrity, capillary stability, vascular tone, extracellular matrix organization, and endothelial signaling, yet how pericyte subtypes and pericyte-endothelial communication are remodeled across distinct neurodegenerative diseases remains incompletely understood^24^. Addressing this gap can be important for understanding the contribution of the brain vasculature to neurodegeneration and for identifying vascular pathways with diagnostic or therapeutic potential.

Here, we performed a comparative molecular analysis of human brain vascular and perivascular cell types across AD^21^, FTD^22^, and HD^23^ by integrating single-cell and single-nucleus RNA sequencing datasets from the human brain vasculature. This enabled direct cross-disease comparison and systematic identification of shared and disease-specific transcriptional programs across vascular populations. We focused on endothelial zonation^20^, pericyte subtype remodeling^21^, disease-associated gene expression, pathway enrichment, and pericyte-endothelial ligand-receptor communication. Across diseases, we identify both convergent and disease-specific vascular signatures, with prominent remodeling of extracellular matrix, adhesion, basement membrane, and neurovascular signaling pathways.

## Results

### A cross-disease single-nucleus transcriptomic atlas of blood-brain barrier cells

To systematically compare vascular and glial cell populations across neurodegenerative diseases, we reprocessed and harmonized three published brain single-nucleus/single-cell RNA-seq datasets from human AD cortex^21^, FTD-GRN frontal and temporal cortex^22^, and HD neostriatum^23^. To reduce anatomical heterogeneity, we restricted the analysis to disease-relevant brain regions present in each dataset: cortex for AD, frontal and temporal cortical regions for FTD-GRN, and neostriatum for HD. After dataset-specific quality control, normalization, and dimensionality reduction, the final harmonized analysis included 97,316 nuclei from the AD cortex dataset, 429,435 from the FTD-GRN cortical dataset, and 47,369 from the HD neostriatal dataset. Major vascular, perivascular, glial, and immune populations were annotated and harmonized across datasets using established marker genes. We concentrated our analysis on endothelial cells (*CLDN5*, *PECAM1*, *VWF*), pericytes (*PDGFRB*, *RGS5*), smooth muscle cells (SMCs) (*ACTA2*, *TAGLN*), fibroblasts (*COL1A1*, *COL1A2*, *DCN*), astrocytes (*AQP4*, *GFAP*, *SLC1A2*), oligodendrocytes (*MBP*, *PLP1*, *MOG*), microglia (*P2RY12*, *TMEM119*, *CX3CR1*), and macrophages (*CD163*, *MRC1*, *LYZ*) (Supp. Fig. 1A). UMAP visualization revealed consistent clustering of major vascular and glial populations across cohorts, indicating successful integration and annotation of shared cell types (Fig. 1A). To evaluate dataset compatibility, we calculated pairwise Pearson correlations between matched control populations from each disease cohort. Control transcriptomes exhibited strong concordance across datasets, with most cell-type comparisons showing high correlation coefficients (r > 0.85), demonstrating that transcriptional identities were preserved across studies and supporting subsequent cross-disease comparisons (Fig. 1B). Despite differences in total cell numbers between studies, besides macrophages, all major cell populations were represented across datasets (Fig. 1B,C). We next compared the relative abundance of major cell populations between control and disease samples across the three neurodegenerative disorders. Across all diseases, endothelial cells were proportionally reduced in disease compared with controls (AD: 37.1% vs. 17.2%; FTD-GRN: 2.3% vs. 1.2%; HD: 5.8% vs. 3.4%), whereas oligodendrocytes showed a relative increase (AD: 10.9% vs. 42.3%; FTD-GRN: 1.4% vs. 2.2%; HD: 51.0% vs. 57.9%). The most pronounced changes were observed in AD, which additionally exhibited a marked reduction in pericytes (29.0% vs. 14.7%) and an expansion of astrocytes (10.9% vs. 17.9%). In contrast, HD displayed the smallest alterations in cellular composition between control and disease samples, indicating comparatively limited changes in cell-type abundance despite widespread transcriptional remodeling (Fig. 1C).

**Figure 1:**
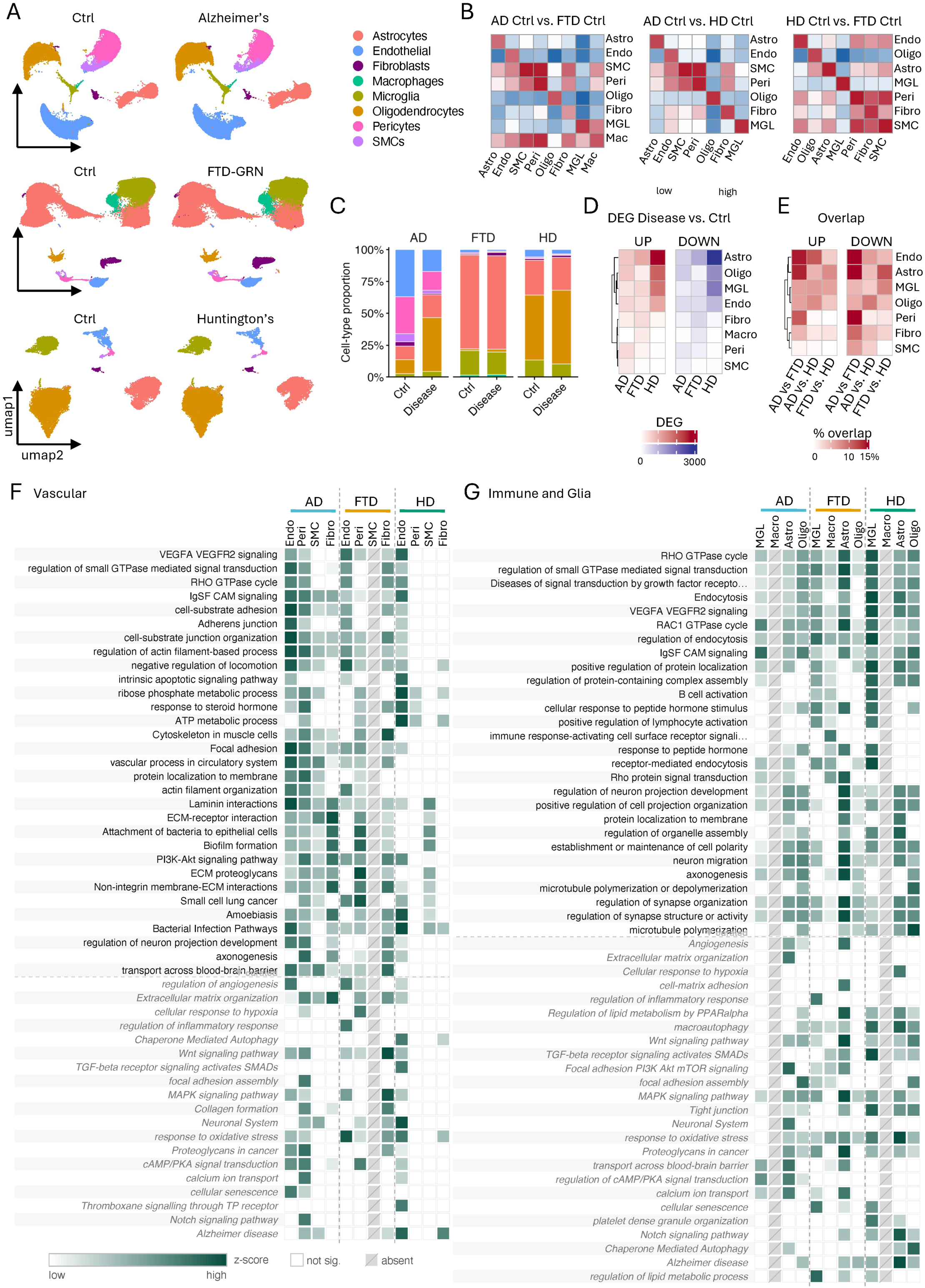
Cross-disease single-nucleus transcriptomic atlas of brain vascular and glial cells. **A**, UMAP visualization of harmonized non-neuronal nuclei from AD cortex, FTD-GRN frontal/temporal cortex, and HD neostriatum, split by condition (control versus disease) and colored by cell type. Control and disease samples are shown separately for each cohort. **B,** Pairwise Pearson correlation heatmaps of cell-type-level transcriptomes between control samples across datasets (AD Ctrl vs. FTD Ctrl, AD Ctrl vs. HD Ctrl, HD Ctrl vs. FTD Ctrl). Color scale indicates correlation coefficient. **C**, Cell-type composition across control and disease samples for each cohort. Stacked bars show the relative abundance of annotated cell classes within AD, FTD-GRN, and HD datasets. **D**, Summary of disease-associated transcriptional changes by cell type. Heatmaps show the number of significantly upregulated or downregulated genes (MAST, adjusted p < 0.05, |log2FC| > 0.25) in disease versus control for each major cell class and disease cohort. **E**, Cross-disease overlap of disease-associated differentially expressed genes. Heatmaps show the percentage overlap of upregulated and downregulated genes between disease pairs across major cell classes. **F,G**, Pathway-enrichment overview of vascular cell populations (F) and microglia/macrophages, astrocytes, and oligodendrocytes (G) across AD, FTD-GRN, and HD. Heatmaps summarize enriched pathways in endothelial cells, pericytes, SMCs, and fibroblasts ranked by -log10(padj), also showing curated, disease-associated pathways in grey. Color intensity in pathway heatmaps indicates relative enrichment z-score; white boxes indicate non-significant terms and grey diagonal boxes indicate absent comparisons. AD - Alzheimer’s disease, FTD-GRN - frontotemporal dementia caused by GRN mutation HD - Huntington’s disease.

### Differential expression analysis reveals cell-type-specific and disease-dependent transcriptomic changes

Comparison of disease-associated gene signatures revealed only limited overlap of individual DEGs between AD, FTD-GRN, and HD, indicating that most transcriptional responses are disease-specific despite affecting similar cell types (Fig. 1E).

Having established dataset comparability, we next examined disease-associated transcriptional alterations within each cell type using MAST (adjusted p < 0.05, |log2FC| > 0.25). Differential expression analysis identified substantial disease-induced transcriptional remodeling across both vascular and glial compartments, with oligodendrocytes displaying the largest number of differentially expressed genes (DEGs) in all three diseases, followed by astrocytes, microglia, and vascular populations (Fig. 1D, Supp. Fig. 1B). HD showed the most pronounced transcriptional changes overall, with astrocytes, microglia, and oligodendrocytes each exhibiting more than 1,000 DEGs.

In contrast, AD-associated transcriptional changes were particularly enriched in vascular populations, with pericytes and SMCs displaying substantial numbers of DEGs, whereas these cell types were comparatively less affected in HD and FTD-GRN. FTD-GRN exhibited a distinct pattern, characterized by prominent fibroblast and macrophage responses, which were largely absent in the other diseases.

Comparison of disease-associated DEGs revealed both shared and disease-specific transcriptional signatures (Fig. 1E, Supp. Fig. 1C). Among the most consistently upregulated genes across all three diseases were CNTNAP4, MOBP, and PI16, while ADAMTS9, DLGAP2, and SLCO4A1 were consistently downregulated (Supp. Fig. 1B). A core set of 49 genes was dysregulated across all eight cell types and all three diseases, suggesting a conserved transcriptional response to neurodegeneration that is largely independent of cell identity. Despite this shared signature, the extent of overlap varied substantially between cell types. Endothelial cells, astrocytes, and pericytes exhibited the highest cross-disease concordance, with AD and FTD-GRN showing the greatest DEG overlap. In contrast, HD contributed the largest proportion of disease-specific DEGs across most cell populations, highlighting the distinct transcriptional landscape associated with Huntington’s disease (Fig. 1E, Supp. Fig. 1C).

To identify functionally altered programs in neurodegeneration, we performed gene set enrichment analysis (fgsea) using GO (BP: Biological Process, MF: Molecular Function, CC: Cellular Component), REACTOME, KEGG, and WikiPathways. Of 101,810 pathway-cell type tests, 6,132 (6.0%) were significantly enriched (adjusted p < 0.05), indicating extensive pathway-level remodeling despite limited overlap of individual DEGs across diseases. In vascular cells, hierarchical clustering of normalized enrichment scores (NES) revealed convergent activation of VEGFA-VEGFR2 signaling, small GTPase-mediated signal transduction, and cell-substrate adhesion across diseases, especially visible across vascular compartments, supporting altered endothelial and vascular growth/signaling programs. (Fig. 1F). Pathways associated with BBB transport and angiogenesis were particularly enriched in AD endothelial cells and pericytes, consistent with prominent vascular dysfunction in AD. Extracellular matrix organization, laminin interactions, and collagen formation were enriched in fibroblasts from both AD and FTD-GRN, whereas Notch signaling was selectively activated in AD, and Wnt and TGF-`β` receptor signaling were preferentially enriched in FTD-GRN vascular populations.

In glial and immune cells, a distinct set of pathways emerged (Fig. 1G). Microglia and macrophages from FTD-GRN showed strong enrichment of immune activation programs, including B cell activation, lymphocyte activation, and immune receptor signaling, consistent with the established role of GRN haploinsufficiency in driving dysregulated innate immune responses. HD displayed a prominent translational signature, with ribosomal biogenesis and protein synthesis pathways among the most strongly upregulated programs across the entire dataset (NES = 2.91 in HD endothelial cells). By contrast, FTD microglia showed the strongest downregulation of translational machinery (NES = -3.13 for cytoplasmic ribosomal proteins), indicating opposing regulation of protein synthesis across diseases. Despite these differences, several pathways were consistently altered across diseases and cell types, including chaperone-mediated autophagy, oxidative stress responses, hypoxia and neuronal system-associated signaling, as well as pathways involved in protein quality control. These shared signatures suggest that distinct neurodegenerative diseases converge on common cellular stress-response mechanisms and neurovascular dysfunction programs despite engaging largely disease-specific transcriptional networks.

### Endothelial remodeling converges on capillary vascular segments across neurodegenerative diseases

Given the consistent involvement of endothelial cells across diseases, we next investigated whether these disease-associated changes were distributed uniformly across the arteriovenous axis, along which endothelial cells differ in barrier properties, transporter expression profiles, and immune functions. To address this, endothelial cells were subclustered into arterial, capillary, and venous populations using established zonation markers (Fig. 2A; Supp. Fig. 2A). Capillary endothelial cells constituted the dominant endothelial population across all conditions. In AD, endothelial cell numbers were markedly reduced across all vascular segments, with the most pronounced decrease observed in capillary endothelial cells (∼43% reduction), whereas endothelial populations remained comparatively stable in FTD-GRN and HD (Fig. 2B, Supp. Fig. 2B).

**Figure 2:**
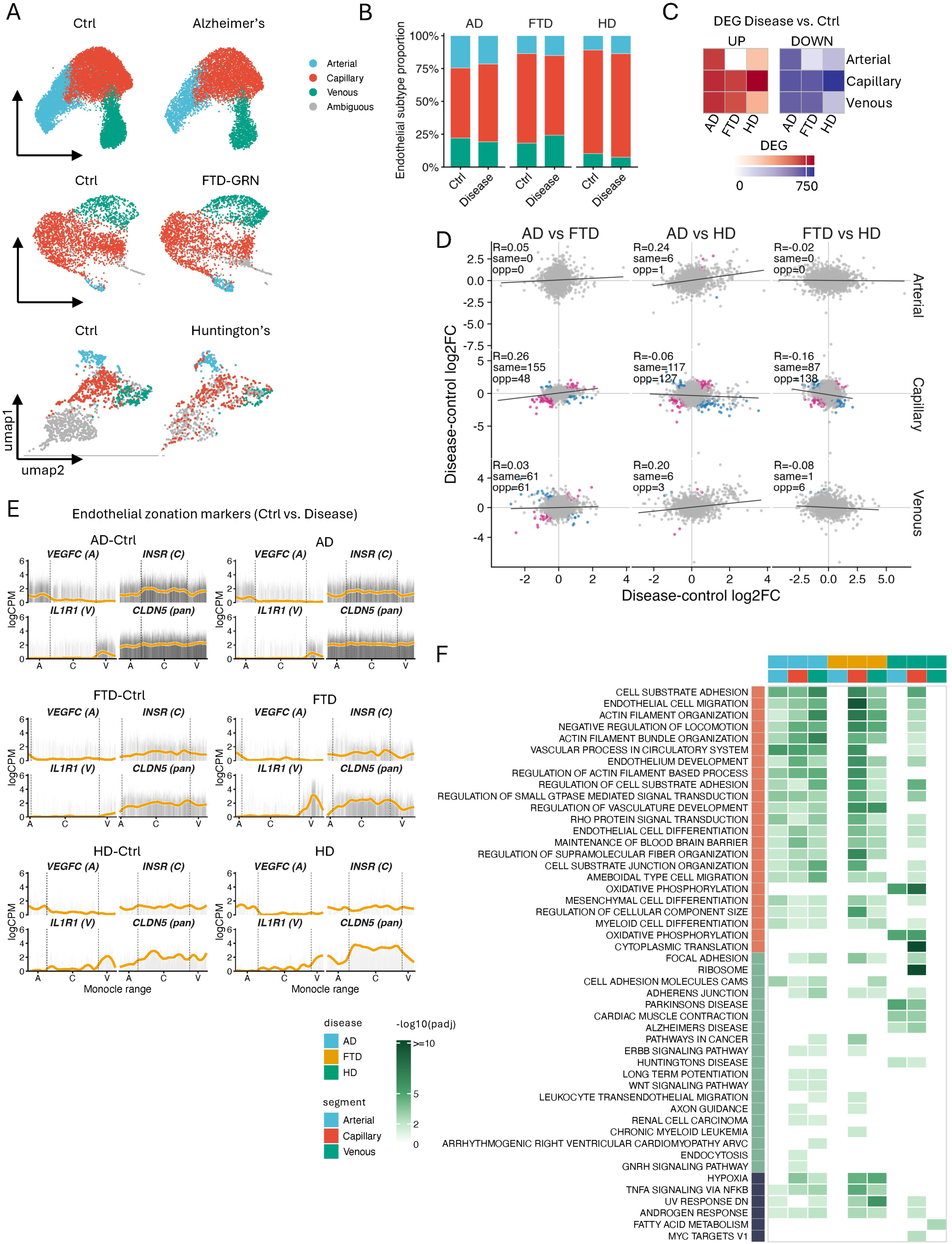
Endothelial zonation analysis across neurodegenerative diseases. **A**, UMAP embeddings of endothelial cells colored by vascular segment (arterial, capillary, venous), split by condition (control versus disease) for AD, FTD-GRN, and HD. **B,** Bar plots of endothelial cell numbers per vascular segment in control versus disease for each cohort. **C,** Heatmap of the number of significant DEGs per endothelial segment and disease, separated by direction (upregulated and downregulated). **D.** Cross-disease correlation of endothelial disease signatures. Scatterplots compare disease-versus-control log2FC values between AD, FTD-GRN, and HD within arterial, capillary, and venous endothelial cells. Pearson’s R, shared directionality, and overlap significance are shown. **E.** Endothelial zonation marker expression across the arterial–capillary–venous axis in control and disease. Smoothed profiles show segment-associated markers and preservation or disease-associated shifts of zonation patterns. **F,** Pathway over-representation analysis of endothelial segments. Rows indicate enriched terms; columns are grouped by disease and colored by vascular segment (arterial, red; capillary, orange; venous, blue). Color intensity represents -log10(adjusted p). Source database indicated by colored sidebar (GO_BP, KEGG, REACTOME). AD - Alzheimer’s disease, FTD-GRN - frontotemporal dementia caused by GRN mutation, HD - Huntington’s disease.

To determine whether endothelial dysfunction was segment-specific, we performed differential expression analysis within each vascular compartment. Capillary and venous endothelial cells carried most transcriptional changes, whereas arterial endothelial cells were comparatively less affected (Fig. 2C). AD exhibited the strongest transcriptional remodeling in capillary endothelial cells, while HD showed the largest response in venous endothelial cells. In contrast, FTD-GRN was associated with relatively modest transcriptional alterations across all endothelial segments.

To assess the extent to which disease-associated endothelial responses were shared across neurodegenerative disorders, we compared differential expression signatures between disease pairs within each vascular segment. Capillary endothelial cells exhibited the greatest cross-disease concordance, particularly between AD and FTD-GRN (r = 0.26), whereas arterial and venous endothelial cells showed little correlation between diseases (Fig. 2D). Comparisons involving HD generally displayed weaker concordance, indicating a more distinct endothelial transcriptional response.

Examination of segment-specific DEGs further highlighted both shared and disease-specific molecular alterations (Supp. Fig. 2C). In AD, endothelial remodeling was characterized by differential expression of genes involved in vascular homeostasis and BBB function, including downregulation of *VEGFC*, *FLT1*, and *PECAM1*, together with increased expression of *KLF2*, *COL4A2*, and *MFSD2A*. FTD-GRN endothelial cells exhibited altered expression of vascular signaling and inflammation-associated genes, including *PICALM*, *TEK*, and *IL1R1*. By contrast, HD endothelial cells showed prominent induction of *CLU*, *TIMP1*, and *NR2F2*, as well as several neurodegeneration-associated genes, including *BIN1*, *GRN*, *OPTN*, and *SORL1*. Together, these findings indicate that distinct neurodegenerative diseases converge on endothelial dysfunction while engaging partially distinct molecular programs.

To examine the impact of neurodegeneration on endothelial zonation, we reconstructed the arteriovenous trajectory and assessed the expression of canonical zonation markers. The arterial marker *VEGFC*, capillary marker *INSR*, and venous marker *IL1R1* retained their expected spatial expression patterns across diseases, indicating preservation of overall endothelial identity (Fig. 2E). However, disease-associated alterations in expression dynamics were evident, particularly within capillary and venous regions, suggesting that neurodegeneration remodels endothelial gene expression while maintaining the overall zonation architecture.

Pathway enrichment analysis revealed both shared and segment-specific responses along the arteriovenous axis (Fig. 2F). Capillary endothelial cells displayed the broadest pathway alterations across all three diseases, identifying the capillary bed as the principal site of endothelial remodeling. The most consistently enriched pathways were related to cytoskeletal organization and barrier function, including actin filament organization, cell-substrate adhesion, endothelial cell migration, regulation of small GTPase signaling, and maintenance of BBB integrity. These programs were particularly prominent in AD and HD, suggesting widespread remodeling of endothelial architecture and vascular homeostasis. Venous endothelial cells exhibited a partially overlapping response characterized by enrichment of adherens junction, focal adhesion, and VEGF signaling pathways, indicating shared regulation of endothelial–matrix interactions across vascular segments. In contrast, disease-specific programs were more restricted. HD endothelial cells showed prominent enrichment of oxidative phosphorylation, cytoplasmic translation, and ribosomal pathways, consistent with extensive metabolic and translational remodeling. FTD-GRN endothelial cells preferentially enriched immune-related pathways, including antigen processing and presentation, whereas AD exhibited the strongest enrichment of pathways linked to BBB maintenance and endothelial migration. Across all diseases, endothelial stress-response pathways, including hypoxia and TNFα signaling *via* NFκB, were recurrently enriched, suggesting that diverse neurodegenerative disorders converge on common vascular adaptation programs despite engaging distinct disease-specific molecular networks. Together, these findings identify capillary endothelial cells as a major site of convergent vascular dysfunction across neurodegenerative diseases.

### Pericyte sub-clustering reveals selective loss of matrix-type pericytes across diseases

Subclustering of pericyte nuclei across all three datasets identified two transcriptionally distinct pericyte states that exhibited differential vulnerability to neurodegeneration (Fig. 3A,B, Supp. Fig. 3A). One state was enriched for extracellular matrix and vascular remodeling genes, including *COL4A1*, *COL4A2*, *ADAMTS1*, and *ADAMTS9*, whereas the other preferentially expressed transport- and signaling-associated genes such as *SLC6A1*, *SLC6A13*, *APOD*, and *SLC20A2* (Fig. 3C; Supp. Fig. 3B). These populations correspond to the previously described mesh pericyte (M-peri) and tip pericyte (T-peri) states, respectively^21^. Across all three diseases, M-peri proportion declined relative to matched controls (Fig. 3B; Supp. Fig. 3A). In AD and FTD controls, M-peri constituted a minority of total pericytes (∼27% and ∼26%, respectively). In HD controls, the composition was markedly different, with M-peri representing the majority subtype (∼55%), suggesting a baseline difference in pericyte subtype composition between cortical and striatal vasculature. Nevertheless, disease was associated with a pronounced reduction of M-peri cells in all three disorders (AD: 26.7% to 6.7%; FTD-GRN: 25.9% to 9.8%; HD: 54.9% to 33.5%), resulting in a relative shift toward T-peri predominance. Despite differences in baseline abundance, selective depletion of M-peri emerged as a shared feature of neurodegeneration. Overlap analysis of DEGs (adjusted p < 0.05, |log2FC| > 0.5) identified a core set of genes dysregulated across all three diseases, including *COL4A2*, *MIR4435-2HG*, *OSMR*, *PDE10A*, *PCBP3*, and *TRPC4* (Fig. 3D). Consistent with this classification, canonical pericyte markers were retained across both subtypes, while subtype-specific gene expression patterns were preserved across diseases, indicating that disease-associated changes reflect remodeling of established pericyte states rather than the emergence of novel populations (Supp. Fig. 3B). To further assess the extent of transcriptional convergence, we compared differential expression signatures between disease pairs within each pericyte subtype. M-peri cells exhibited greater cross-disease concordance than T-peri cells, with the strongest similarity observed between AD and FTD-GRN (r = 0.31) (Fig. 3E). Comparisons involving HD showed weaker correlations, consistent with the data from the endothelial subpopulations, indicating a more distinct pericyte response in the striatum. Together, these findings identify matrix-associated pericytes as a major site of convergent transcriptional remodeling across neurodegenerative diseases. To further characterize the shared and specific transcriptional programs underlying pericyte remodeling, we clustered genes that were differentially expressed across the three diseases into co-regulated modules (Fig. 3F). These modules revealed coordinated alterations in pathways related to extracellular matrix organization, vascular signaling, hypoxia responses, endothelial migration, and blood`–`brain barrier regulation. While individual genes exhibited disease-specific patterns, many module-level responses were consistently altered across AD, FTD-GRN, and HD, indicating convergence on common neurovascular stress and remodeling programs. Notably, extracellular matrix-associated genes and vascular remodeling factors displayed the most consistent cross-disease regulation, supporting the notion that disruption of pericyte-mediated vascular support represents a shared feature of neurodegeneration. Gene set enrichment analysis revealed marked differences in pathway-level responses across diseases and highlighted subtype-specific pericyte programs (Fig. 3G). Hallmark enrichment analysis indicated a shift between structural remodeling pathways and inflammatory-metabolic signaling. Pathways enriched among upregulated genes, including epithelial–mesenchymal transition, apical junction organization, and myogenesis, were preferentially associated with T-peri-enriched transcriptional programs. In contrast, pathways enriched among downregulated genes, including hypoxia, TNFα–NFκB signaling, IL6–JAK–STAT signaling, mTORC1 signaling, Notch signaling, interferon responses, and angiogenesis, showed greater overlap with M-peri-associated genes, suggesting preferential suppression of matrix-associated pericyte functions in disease.

**Figure 3.**
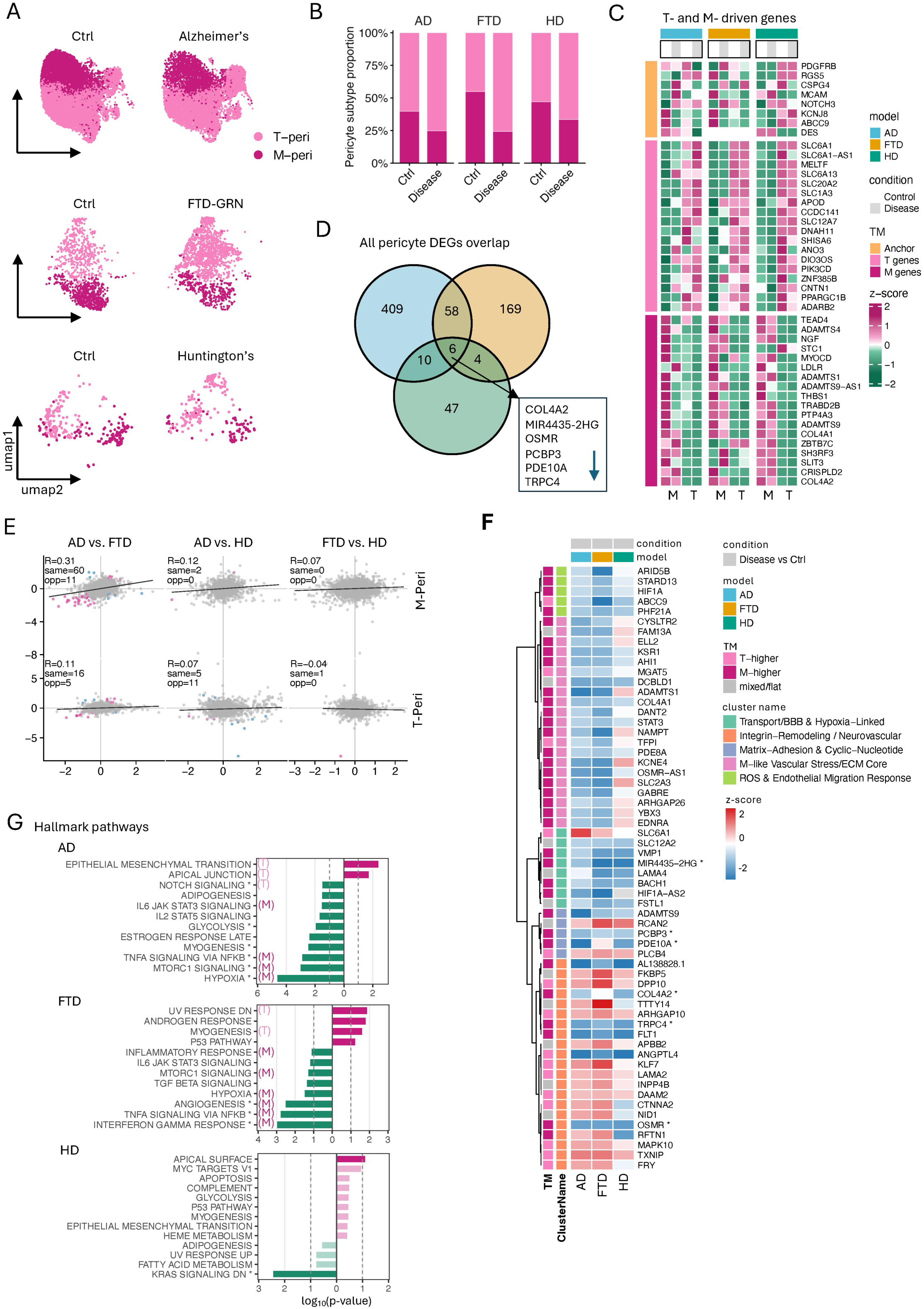
Matrix-associated pericytes are selectively depleted and transcriptionally remodeled across neurodegenerative diseases. **A**, UMAP visualization of pericyte subclusters across AD, FTD-GRN, and HD datasets, identifying mesh pericytes (M-peri) and tip pericytes (T-peri). **B**, Relative abundance of M-peri and T-peri populations in control and disease samples. Numbers indicate the proportion of each subtype within the pericyte compartment. **C**, Heatmap showing expression changes of genes associated with T- and M-peri subpopulations. Rows represent DEGs and columns represent disease conditions. Colors indicate scaled log2 fold change relative to controls. **D**, Venn diagrams showing overlap of significantly differentially expressed genes (DEGs) across AD, FTD-GRN, and HD. Shared genes across all three diseases are highlighted. **E**, Pairwise comparison of disease-associated transcriptional responses between pericyte subtypes. Scatter plots show log2 fold changes for shared genes between disease pairs. Pearson correlation coefficients (R) and the number of concordantly or discordantly regulated genes are indicated. **F**, Heatmap of shared pericyte DEGs grouped into co-regulated functional modules. Gene modules are annotated according to major biological processes, including extracellular matrix remodeling, vascular signaling, hypoxia responses, endothelial migration, and neurovascular stress pathways. Colors represent scaled gene expression changes across diseases. Asterisks indicate the 6 genes being consistently regulated across diseases**. G**, Hallmark pathway enrichment analysis of disease-associated pericyte transcriptional changes (MAST was performed separately on T- and M-peri). Bars indicate significantly enriched pathways among upregulated (right) and downregulated (left) genes for each disease. Asterisks indicate curated pathways. (T) and (M) indicate enriched terms whose contributing genes preferentially overlapped with T-peri- or M-peri-enriched signatures, respectively.

AD exhibited the most extensive pathway remodeling, with significant alterations across both T-peri- and M-peri-associated programs. FTD-GRN displayed a more restricted but partially overlapping profile characterized by changes in angiogenesis, inflammatory signaling, hypoxia responses, and p53-associated stress pathways. In contrast, HD showed comparatively limited pathway enrichment, consistent with the small number of subtype-specific DEGs identified in HD pericytes.

GO enrichment analysis further supported a subtype-skewed pericyte response (Supp. Fig. 3C). Upregulated terms related to neuronal projection development, synaptic organization, cytoskeletal remodeling, and actin- and tubulin-binding preferentially overlapped with T-peri-enriched genes. Conversely, downregulated pathways involving hypoxia responses, extracellular matrix organization, collagen-containing structures, basement membrane components, cAMP/PKA signaling, and phosphodiesterase activity were more strongly associated with M-peri-enriched genes. Consistent with these findings, pooled pathway enrichment analysis identified broad alterations in cell communication, intracellular signaling, extracellular matrix organization, focal adhesion, HIF-1 signaling, PDGF signaling, and Rho GTPase-mediated pathways, with the strongest enrichment observed in AD, weaker enrichment in FTD-GRN, and little detectable pathway-level remodeling in HD (Supp. Fig. 3D). Together, these findings indicate that while selective depletion of matrix-associated pericytes is shared across neurodegenerative diseases, the accompanying transcriptional remodeling is dominated by suppression of M-peri-associated programs and is most pronounced in AD.

### Conserved and disease-specific remodeling of endothelial–pericyte communication across neurodegenerative diseases

Given the convergent transcriptional remodeling observed in endothelial cells and pericytes across neurodegenerative diseases, we next investigated how these changes affect intercellular communication within the neurovascular unit. Using CellChat, we reconstructed ligand–receptor interactions between endothelial cells and pericytes in control and disease samples from AD, FTD-GRN, and HD (Fig. 4, Supp. Fig. 4). We focused on this axis because endothelial–pericyte crosstalk maintains BBB integrity, vessel stability, and trophic support, and because both partners showed consistent transcriptomic perturbation across AD, FTD-GRN, and HD (Fig. 1-3).

**Figure 4.**
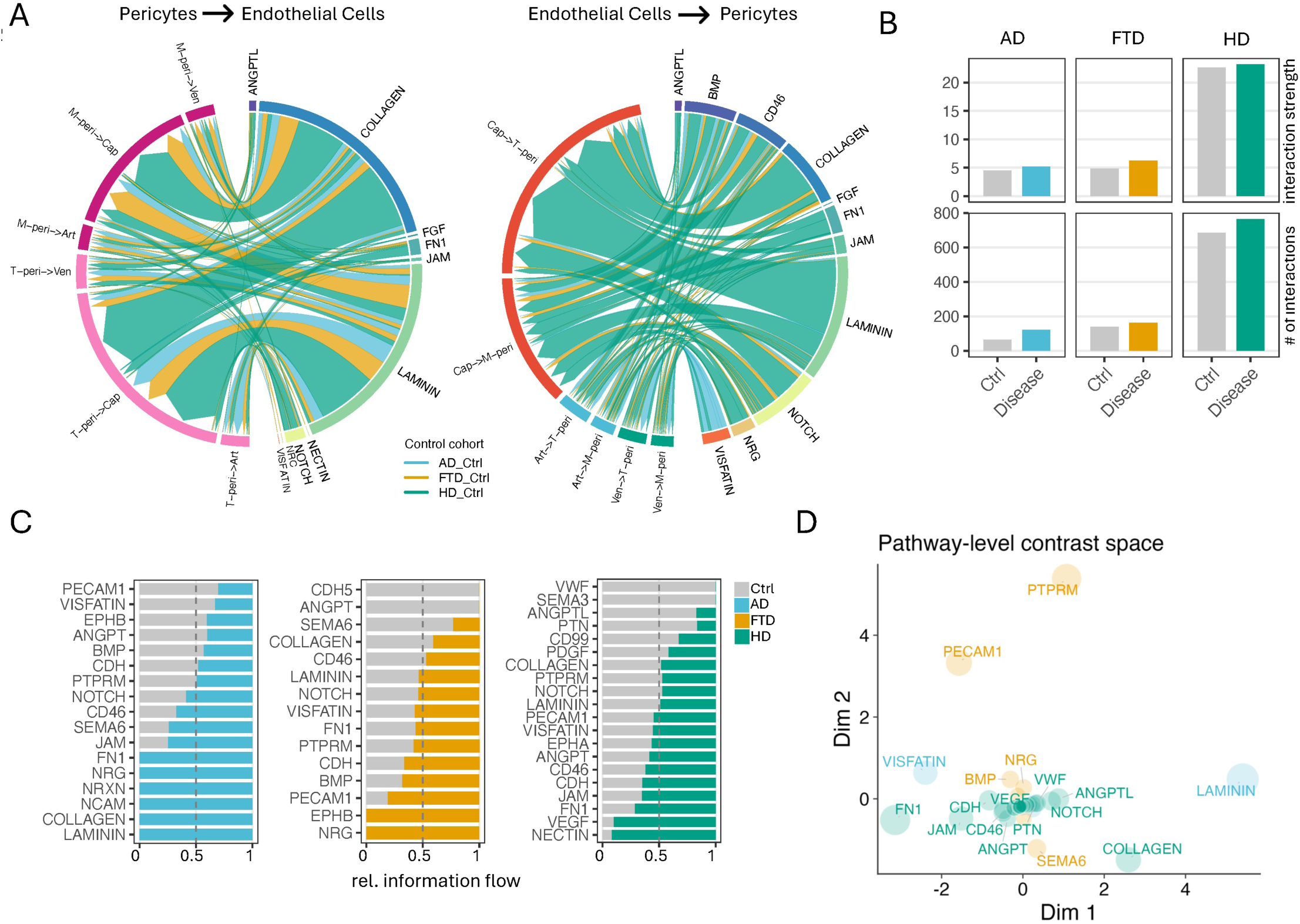
CellChat analysis of pericyte–endothelial signaling across AD, FTD-GRN, and HD. **A**. Shared signaling pathways in control samples. Thirteen CellChat signaling pathways were identified exclusively in control networks and were shared across all three disease cohorts. **B**. Global interaction density of pericyte–endothelial CellChat networks. Bar plots show the total number of inferred CellChat interactions and the cumulative interaction strength between pericyte and endothelial cell populations in control and disease samples for each cohort. Grey bars represent control samples, whereas disease samples are colored according to disease group: AD (blue), FTD-GRN (orange), and HD (green). Interaction number corresponds to the count of ligand–receptor interactions passing CellChat filtering criteria, while total interaction strength represents the summed communication weight across the inferred network. **C**. Relative pathway information flow in disease-specific CellChat networks. Stacked bar plots display the relative contribution of selected signaling pathways to total information flow in control and disease networks for AD, FTD-GRN, and HD. For each cohort, pathway information flow was normalized between the corresponding control and disease conditions. Grey indicates control, while disease-specific colors indicate disease samples: AD (blue), FTD-GRN (orange), and HD (green). The dashed vertical line marks equal contribution from control and disease conditions. **D**. Pathway-level contrast space of disease-associated CellChat changes. Signaling pathways were projected into a two-dimensional contrast space according to their Disease–Control differences in information flow across AD, FTD-GRN, and HD. Pathways positioned close together exhibit similar patterns of change across diseases, whereas distant pathways represent disease-specific signaling alterations. Pathway labels are colored according to the disease comparison showing the largest absolute change: AD (blue), FTD-GRN (orange), and HD (green). Background circles indicate the magnitude of the strongest Disease–Control change for each pathway.

We first used a sensitivity-oriented CellChat analysis to survey the broad repertoire of endothelial-pericyte signaling pathways present in control tissues (Fig. 4A). This analysis used 10% truncated mean and population-size weighting to retain sparse extracellular-matrix-associated interaction. In control tissue, communication between endothelial cells and pericytes was dominated by extracellular matrix-associated and contact-dependent signaling pathways, including LAMININ, COLLAGEN, FN1, JAM, NOTCH, and ANGPTL signaling. Endothelial-to-pericyte communication additionally involved trophic and metabolic pathways such as BMP, VISFATIN, NRG, and CD46 signaling, consistent with established roles in vascular maintenance, maturation, and BBB homeostasis (Fig. 4A). We then performed the main disease-control comparison using a conservative CellChat workflow based on the triMean estimator and population.size = FALSE, allowing us to compare inferred per-cell signaling strength independently of cell abundance (Fig. 4B-D). Disease induced extensive remodeling of the endothelial-pericyte communication network. Surprisingly, the overall number and strength of inferred interactions were maintained or modestly increased across all three diseases (Fig. 4B), indicating that neurodegeneration does not simply lead to loss of per-cell vascular communication, but rather to network rewiring.

Pathway-level analysis revealed both shared and disease-specific alterations (Fig. 4C, Supp, Fig. 4A). Across diseases, extracellular matrix-associated and adhesion pathways, including LAMININ, COLLAGEN, FN1, and NCAM signaling, showed prominent disease-associated changes, although the direction and magnitude of these changes differed across cohorts. Additional disease-specific changes were observed, including preferential reduction of BMP, EPHB, and PECAM1 signaling in FTD-GRN and attenuation of VEGF-and NECTIN-associated pathways in HD. To compare signaling programs across diseases, we projected pathway-level communication changes into a shared contrast space (Fig. 4D). Pathways located near the center of this projection showed broadly similar or relatively modest shifts across AD, FTD-GRN, and HD, including core vascular signaling pathways such as ANGPT, JAM, CD46, PTN, VEGF, NOTCH, and ANGPTL. In contrast, pathways positioned further from the center represented stronger or more disease-selective remodeling. ECM-associated pathways contributed prominently to this separation, with LAMININ showing a stronger AD-associated shift and COLLAGEN showing the largest absolute shift in HD, while PTPRM and PECAM1 were more strongly associated with FTD-GRN-specific changes.

To ensure that these findings were not driven by disease-associated differences in cell abundance, we repeated the CellChat analysis after balancing endothelial and pericyte cell numbers across conditions. The balanced analysis confirmed robust extracellular matrix-associated signaling changes, with COLLAGEN emerging as enriched also in FTD-GRN and LAMININ remaining detectable across disease comparisons. Additional pathways, including FN1, PECAM1, CD46, PTPRM, and SEMA6, were also retained, supporting the interpretation that altered communication primarily reflects intercellular remodeling rather than cell-composition effects (Supp. Fig. 4B). Sender`–`receiver analysis revealed disease-specific redistribution of signaling flow between endothelial and pericyte populations (Supp. Fig. 4C). In AD, the largest increases were observed in signaling originating from both M-peri and T-peri populations toward capillary and venous endothelial cells, indicating a shift toward pericyte-driven communication. In contrast, FTD-GRN was characterized by enhanced signaling among endothelial populations, particularly from venous endothelial cells, suggesting preferential remodeling of endothelial-centered communication networks. HD displayed a more heterogeneous pattern involving both endothelial and pericyte populations, consistent with the extensive network rewiring observed throughout the HD neurovascular compartment. These findings indicate that although common signaling pathways are affected across diseases, the cellular architecture through which these signals are transmitted differs substantially between disease contexts.

## Discussion

The extent to which major neurodegenerative diseases converge or diverge at the level of the brain vasculature has remained largely unexplored. Here, we address this gap by integrating publicly available vessel-enriched human snRNA-seq datasets across AD, FTD-GRN, and HD to build a molecular atlas of endothelial cells and pericytes across the diseases^21–23^. Our analysis reveals a conserved vascular response to neurodegeneration and disease-specific transcriptional programs that may reflect the distinct pathological pathways of each neurodegenerative disease along the arteriole-to-venule axis.

We identified a conserved transcriptional core shared across all three diseases and all eight cell types. This set of 49 genes is dysregulated regardless of cell identity or disease context. This "pan-neurodegenerative" signature, which includes consistent upregulation of *CNTNAP4* and *MOBP* and downregulation of *ADAMTS9*, *DLGAP2*, and *SLCO4A1*, suggests a generalized molecular response to neurodegeneration, irrespective of the primary proteinopathy. Beyond transcriptional changes within cell types, we observe compositional shifts in the vascular compartment across all three diseases, with the most pronounced in AD. The proportional loss of endothelial cells ranges from 37.1% to 17.2% in AD. HD and FTD-GRN show a more modest loss of endothelial cells. AD vascular density changes and capillary loss have been reported in several studies^25,26^. Whereas for HD and FTD-GRN, vascular density changes and capillary loss have not been investigated. HD showed the smallest compositional shifts between disease and control, which may be due to the brain regions analyzed, as HD was sourced from striatal regions, whereas AD and FTD-GRN were sourced from cortical regions.

Analysis of the vasculature along the arteriole-venule axis revealed disease-specific alterations in vessel subtypes^20,21^. Overall, pathways related to actin dynamics, cell-substrate adhesion, endothelial migration, and BBB maintenance are upregulated in most vessel subtypes across all three diseases, particularly in capillaries and venules. Arterioles from AD and HD also share these pathways. The FTD-GRN dataset had too few arterioles to include them in the analysis, limiting interpretation. Alongside these shared vascular changes, we identify disease-specific transcriptional programs that likely reflect distinct pathological environments of each disorder. In HD, the arterial endothelial compartment showed enrichment in platelet activation signaling and RHO-GTPase cycling, suggesting a pro-thrombotic arteriole-platelet interaction and a cytoskeletal remodeling response. In FTD-GRN, CAM adhesion and ECM remodeling were specifically altered, suggesting immune cell-vascular interaction in the venules’ endothelial cells^27^. AD shows similar pathways related to immune cell-vascular interaction altered in capillaries and venules, a phenomenon that has been described, contributing to reduced blood flow and cognitive impairment in mouse models^28^. These disease-specific endothelial signatures suggest that all three diseases converge at the capillary level, but that upstream and downstream vascular effectors diverge across diseases, opening the possibility of disease-specific therapeutic interventions.

The analysis identified significant changes in the pericytes in all three disorders. We identified the same two subclusters M-peri and T-peri described in AD also in HD and FTD-GRN, suggesting that pericytes also contribute to vascular dysfunction in HD and FTD-GRN. M-peri are associated with extracellular matrix genes such as laminin, and are linked to cognitive decline, whereas T-peri are not linked to cognitive decline^29^. Generally, across all three diseases, M-peri are consistently reduced relative to controls, despite substantial differences in their baseline proportions. These baseline proportions likely reflect differences between the datasets and, for HD, also the brain region sampled. We found that M-peri constitute a minority subtype in the cortical vasculature: 26.7% in AD, 25.9% in FTD, and 54.9% in HD. The higher number of M-peri in HD likely reflects regional differences, since the sampling occurred in the striatal vasculature. While this limits the interpretation in this work, it underscores the importance of more brain-region-specific sampling to better understand differences in pericyte populations. However, the consistent reduction in M-peri is consistent across these distinct anatomical samples and diseases, suggesting a selective vulnerability of M-peri in the neurodegenerative diseases. This finding is further strengthened by the identification of the ECM genes *COL4A1*, *COL4A2*, and *ADAMTS9* across all three datasets, suggesting a conserved transcriptional program underlying M-peri loss in neurodegenerative diseases.

Nonetheless, within M-peri, differences occurred: for example, in AD and FTD-GRN, M-peri show suppression of angiogenesis, IL6-JAK-STAT3 signaling, TNF-NF-κB signaling, and apoptosis pathways, reflecting a more senescence-like phenotype^30,31^. By contrast, M-peri in HD resulted in activated apoptosis and TNF-NF-κB signaling alongside suppression of mitotic spindle pathways, suggesting an active pro-apoptotic stress response^32,33^. Of course, these changes should be interpreted with caution due to regional sampling differences across the available datasets, but these differences may reflect variations in the underlying disease mechanisms. For example, amyloid and tau pathology in AD and TDP-43 in FTD-GRN may chronically impair pericyte function, the huntingtin aggregation and transcriptional dysregulation characteristic of HD may trigger apoptotic cascades in pericytes. While further work is needed to confirm these results, experimentally, it could suggest that different therapeutic strategies are needed to improve vascular function in these diseases.

The FTD-GRN dataset reveals a vascular and perivascular immune signature that is largely absent from AD and HD, characterized by elevated fibroblast and macrophage responses and strong enrichment of B cell and lymphocyte activation pathways in microglia and macrophages. We have shown that immune cell-vascular interactions are altered in FTD-GRN mice^27^, and two recent studies have reported reduced BBB permeability in FTD, collectively suggesting that neurovascular function is altered in FTD; further investigation is needed to better understand this in detail^34,35^.

Several limitations should be considered when interpreting this analysis. First, the three datasets derive from distinct brain regions, cortical regions in AD and FTD, and the striatum in HD, making it difficult to fully disentangle disease effects from regional differences in the analysis. Future studies are needed; collecting matched regional data across diseases will be essential to resolve region-specific from disease-specific vascular signatures. Second, cross-dataset integration, even with harmony-based batch correction, may incompletely remove technical variation between studies that used different tissue dissociation and library preparation protocols. Third, the cross-sectional nature of these datasets and their interpretation should be taken with caution, as age, disease stage, and other risk factors are not accounted for in this study.

However, overall, this work provides the first integrated cross-disease comparison of the human brain vasculature across these neurodegenerative diseases, offering a rich resource for the field.

## Methods

### Public single-nucleus/single-cell RNA-seq datasets

We re-analysed three previously published human brain single-nucleus/single-cell RNA-seq datasets representing Alzheimer’s disease (AD), progranulin-associated frontotemporal dementia (FTD-GRN), and Huntington’s disease (HD); Supplemental Table 1. To reduce anatomical heterogeneity across disease cohorts, analyses were restricted to disease-relevant brain regions available in each dataset. For AD, only cortical samples were retained. For FTD-GRN, frontal and temporal cortical regions were retained, whereas occipital regions were excluded from the final harmonized analysis. For HD, the analysis was restricted to neostriatal tissue. Control and disease samples were processed separately at the dataset level and then harmonized for cross-disease comparison.

The final harmonized non-neuronal/non-OPC analysis included 97,316 AD nuclei/cells (45,343 control and 51,973 disease), 429,435 FTD-GRN nuclei/cells (135,836 control and 293,599 disease), and 47,369 HD nuclei/cells (22,399 control and 24,970 disease). Final cell classes retained for downstream analysis comprised astrocytes, endothelial cells, fibroblasts, macrophages where present, microglia, oligodendrocytes, pericytes, and smooth muscle cells (SMCs). Neurons, oligodendrocyte precursor cells (OPCs), mesenchymal cells, lymphocytes, and manually identified low-quality or contaminating clusters were excluded from the focused non-neuronal vascular analysis.

### Dataset preprocessing and quality control

All analyses were performed in R using Seurat-based workflows, as previously established.^36–43^ Because the three source datasets differed in file format, acquisition strategy, and original processing, each dataset was first processed with a dataset-specific workflow that followed the quality-control logic of the corresponding publication as closely as possible, while using a common downstream harmonization strategy for cell-type annotation and vascular subsetting.

For the AD dataset, 10x Genomics count matrices from cortical AD and control samples were loaded using ‘Read10X’ and converted into Seurat objects. Initial quality control retained nuclei/cells with at least 200 detected counts, 200-5,000 detected genes, and mitochondrial gene content of at most 5%. Mitochondrial content was calculated using genes matching the ‘^MT-’ prefix. Samples were normalized independently using SCTransform while regressing mitochondrial content. Doublets were removed per sample using DoubletFinder with PCs 1-20, ‘pN = 0.25’, fixed ‘pK = 0.09’, and an expected doublet fraction of 5%, followed by homotypic doublet adjustment. Singlet objects were integrated using Seurat SCT integration with 1,000 integration features. Dimensionality reduction and clustering were performed using PCA, 30 PCs for graph construction and UMAP, and clustering resolution 0.2.

For the FTD-GRN dataset, patient and control samples were loaded per region and converted into Seurat objects using ‘min.cells = 3’ and ‘min.features = 200’. Per-sample QC retained nuclei/cells with 200-6,000 detected genes, no more than 40,000 counts, and mitochondrial content below 5%. Ribosomal gene content was also calculated for QC tracking using ‘^RPL’ and ‘^RPS’ genes. Doublets were removed per sample using scDblFinder. Singlets were merged, log-normalized with a scale factor of 10,000, and variable genes were selected using the vst method with 2,000 highly variable genes. Data were scaled on highly variable genes, PCA was computed, and Harmony batch correction was applied to the first 30 PCs using donor as the grouping variable. The atlas-level graph was constructed from the Harmony embedding using 30 dimensions and ‘k.param = 30’, followed by clustering at resolution 0.1 and UMAP embedding. After major cell-type annotation, only frontal and temporal cortical regions were retained for the final FTD-GRN analysis, and labels were harmonized from ‘DM’/’DM_Ctrl’ to ‘FTD_GRN’/’FTD_GRN_Ctrl’.

For the HD dataset, disease and control count matrices and metadata were loaded from RDS files. Genes expressed in fewer than 50 cells were removed prior to Seurat object construction. QC retained nuclei/cells with 500-8,000 detected genes and mitochondrial content at most 10%. Data were log-normalized using a scale factor of 10,000, 2,000 highly variable genes were selected using the vst method, and scaling was performed with regression of total UMI count and mitochondrial content. PCA was computed using 30 components. UMAP was initially generated from PCA embeddings with ‘min.dist = 0.75’, and Harmony correction was then applied using the available batch-like metadata column, falling back to sample identity when necessary. Clustering was performed on Harmony embeddings using 30 dimensions and resolution 0.2. Doublets were removed with DoubletFinder using a 7.5% expected doublet rate and homotypic adjustment; clustering and UMAP were rerun after doublet removal.

### Cell-type annotation and harmonization

Major cell classes were annotated using canonical marker genes and marker-derived module scores, followed by manual inspection of UMAP embeddings, marker dot plots, and cluster-level marker expression. Marker sets included endothelial markers such as ‘*CLDN5*’, ‘*PECAM1*’, ‘*FLT1*’, and ‘*VWF*’; pericyte markers such as ‘*PDGFRB*’, ‘*RGS5*’, ‘*ABCC9*’, ‘*KCNJ8*’, and ‘*CSPG4*’; SMC markers including ‘*ACTA2*’, ‘*TAGLN*’, ‘*MYH11*’, and ‘*CNN1*’; astrocyte markers including ‘*AQP4*’ and ‘*GFAP*’; oligodendrocyte markers including ‘*PLP1*’, ‘*MOBP*’, and ‘*MBP*’; microglial markers including ‘*P2RY12*’, ‘*CX3CR1*’, and ‘*CSF1R*’; fibroblast markers including ‘*COL1A1*’ and ‘*DCN*’; and macrophage/CAM markers including ‘*F13A1*’ and ‘*MRC1*’. After dataset-level annotation, labels were harmonized across AD, FTD-GRN, and HD to generate a common non-neuronal cell-type framework.

For Figure 1 composition analyses, final harmonized objects were used to calculate cell-type proportions within each disease and condition. Cell-type counts were converted to percentages within each disease-control or disease group separately, so that composition plots reflected relative cell-type abundance within each condition rather than absolute sequencing depth. Endothelial and pericyte subtype proportions were calculated analogously from the focused endothelial and pericyte objects.

### Endothelial cell subclustering and zonation analysis

Endothelial cells were subset from each disease-specific object and analysed separately. Endothelial identities were harmonized into arterial, capillary, and venous segments, with ambiguous cells excluded from the final differential-expression and pathway-enrichment analyses where indicated. Segment assignment was guided by canonical endothelial zonation markers, including arterial-associated ‘*VEGFC*’, capillary-associated ‘*INSR*’, venous-associated ‘*IL1R1*’, and pan-endothelial ‘*CLDN5*’, together with broader marker inspection across endothelial clusters.

To visualize endothelial zonation structure, UMAP embeddings were generated for each disease dataset and endothelial cells were plotted by segment and condition. Zonation marker profiles were additionally visualized along an arterial-to-capillary-to-venous ordering. For these marker-curve plots, endothelial cells assigned to arterial, capillary, or venous states were retained, and Monocle3 was used to learn a graph on the UMAP embedding. Arterial cells were used as root cells to order cells along pseudotime. Cells were then arranged by segment and pseudotime within each condition, and marker expression was displayed across the ordered monocle range. Marker-expression trends were shown using LOESS smoothing (‘span = 0.22’).

### Pericyte subclustering and subtype annotation

Pericytes were subset from each disease-specific object and reclustered to distinguish transcriptionally defined pericyte states. Donors with fewer than 50 pericytes were excluded from subtype reclustering. For FTD-GRN pericytes, cells were split by donor, normalized, and integrated using CCA-based Seurat integration with 3,000 integration features. Integrated pericyte objects were embedded using PCA and UMAP and clustered using graph-based clustering at resolution 0.2. Pericyte subtypes were classified as T-peri or M-peri using module scores derived from marker sets. T-peri markers included ‘*SLC20A2*’, ‘*SLC1A3*’, ‘*SLC6A12*’, ‘*SLC6A1*’, ‘*SLC12A7*’, and ‘*SLC6A13*’, whereas M-peri markers included extracellular matrix-associated genes such as ‘*COL4A1*’, ‘*COL4A2*’, ‘*COL4A3*’, ‘*COL4A4*’, ‘*LAMA4*’, and ‘*ADAMTS1*’. SMC contamination was assessed using markers such as *‘TAGLN*’, ‘*ACTA2*’, ‘*MYH11*’, and ‘*CNN1*’.

### Differential gene-expression analysis

Differential gene expression was performed with MAST using Seurat ‘FindMarkers’, as previously established.^36,37^ For endothelial cells, disease-control comparisons were performed separately within each endothelial segment and disease dataset. Disease cells were compared against matched controls within AD, FTD-GRN, or HD. MAST models included total RNA count (‘nCount_RNA’) as a latent variable, used ‘logfc.threshold = 0’, and required ‘min.pct = 0.10’. Endothelial genes were considered significant for downstream pathway analysis when adjusted P value was below 0.05 and absolute average log2 fold change exceeded 0.25. The final endothelial DE table comprised disease-segment comparisons for AD, FTD-GRN, and HD across arterial, capillary, and venous cells.

For pericytes, disease-control MAST analyses were performed separately within T-peri and M-peri for each disease dataset. MAST models again included ‘nCount_RNA’ as a latent variable, used ‘logfc.threshold = 0’, and required ‘min.pct = 0.10’. Comparisons were skipped if either the disease or control group contained fewer than 30 cells. For pericyte pathway-enrichment analyses, significant genes were defined as adjusted P value below 0.05 and absolute average log2 fold change above 0.5. In sensitivity and cross-disease convergence analyses, a harmonized relaxed threshold of adjusted P value below 0.05 and absolute average log2 fold change above 0.25 was used to match the endothelial analysis and enable direct cross-cell-class comparison.

Region, age, sex, post-mortem interval, and sequencing platform co-vary with disease cohort and were not modeled as covariates in the differential-expression and pathway analyses.

### Pathway-enrichment analysis

For the global analysis, disease-control differential-expression results were generated separately for each harmonized cell class across AD, FTD-GRN, and HD. Differentially expressed genes were identified from MAST results using an adjusted P value below 0.05 and absolute average log2 fold change above 0.25. To assess pathway-level disease responses across the full non-neuronal compartment, pathway enrichment was performed per disease and cell type using MSigDB-derived gene sets. Over-representation analyses were carried out on significant genes using GO Biological Process, KEGG, Reactome, and WikiPathways collections. In parallel, ranked gene-set enrichment analysis was performed using fgsea, with genes ranked by the MAST test statistic where available, or by average log2 fold change otherwise. Hallmark, GO Biological Process, and KEGG gene sets were tested with gene-set sizes restricted to 15-500 genes. Enrichment results were summarized using adjusted P values and normalized enrichment scores to identify shared and cell-type-specific pathway alterations across neurodegenerative diseases. For visualization, pathway heatmaps included both the top-ranked enriched terms and selected biologically relevant pathways of interest, including vascular, extracellular matrix, inflammatory, metabolic, and cell-stress-related pathways. All displayed pathways were derived from the same enrichment outputs; inclusion of selected pathways was used to aid biological interpretation and was not treated as an additional statistical test.

Endothelial pathway enrichment was performed using over-representation analysis on significant MAST-derived genes for each disease and endothelial segment. Gene sets were obtained from MSigDB through ‘msigdbr’ and included GO Biological Process, Hallmark, and KEGG Legacy collections. Enrichment was performed using ‘clusterProfiler::enricher’ with Benjamini-Hochberg correction, ‘pvalueCutoff = 0.05’, ‘qvalueCutoff = 0.2’, ‘minGSSize = 10’, and ‘maxGSSize = 500’. Pathway heatmaps displayed the negative log10 adjusted P value, capped where necessary for visualization.

Pericyte pathway enrichment was performed on subtype- and direction-specific MAST gene sets using g:Profiler. Genes were split by disease, pericyte subtype, and direction of change. Enrichment was run with organism ‘hsapiens’, unordered queries, g:Profiler g:SCS multiple-testing correction, significance threshold 0.05, and sources including GO Biological Process, GO Cellular Component, GO Molecular Function, KEGG, Reactome, Human Phenotype, WikiPathways, and CORUM. Gene sets with fewer than five input genes were not analysed.

### Cross-disease vascular convergence analysis

To quantify shared vascular transcriptional responses across neurodegenerative diseases, disease-control log2 fold changes were compared across AD, FTD-GRN, and HD using the final endothelial and pericyte MAST tables. Endothelial comparisons were performed separately for arterial, capillary, and venous cells, and pericyte comparisons were performed separately for T-peri and M-peri. For each disease-context combination, genes were considered significant when adjusted P value was below 0.05 and absolute log2 fold change exceeded 0.25.

Pairwise disease comparisons were generated for AD versus FTD-GRN, AD versus HD, and FTD-GRN versus HD. For each pair, Pearson and Spearman correlations were calculated across all genes detected in both comparisons. Genes were classified as shared significant, same-direction shared significant, or opposite-direction shared significant based on whether they passed the significance threshold in both diseases and whether the disease-control fold-change direction matched.

Three-disease convergence was defined as genes significant in AD, FTD-GRN, and HD with the same direction of disease-control change in all three comparisons. Additional shared-gene summaries counted genes significant in at least two diseases. Enrichment of shared same-direction gene sets was performed using Hallmark and GO Biological Process collections from MSigDB. Over-representation P values were computed using the hypergeometric test and adjusted within each collection using the Benjamini-Hochberg method.

### Cell-cell communication analysis using CellChat

Cell-cell communication between pericytes and endothelial cells was inferred using CellChat with the human CellChat ligand-receptor database^44,45^ as previously described.^36,37,41,46^ For each disease dataset, pericyte and endothelial objects were merged, and cell groups were defined as T-peri, M-peri, arterial endothelial cells, capillary endothelial cells, and venous endothelial cells. CellChat was run separately for control and disease conditions within each disease dataset. For the main analysis, normalized RNA expression matrices were extracted from the Seurat RNA data layer and supplied to CellChat. Communication probabilities were computed using the conservative triMean estimator, and population.size = FALSE was used so that inferred communication reflected per-cell signaling strength rather than cell abundance-weighted signaling. Interactions were filtered with min.cells = 15. For each condition, overexpressed genes and interactions were identified, ligand-receptor communication probabilities were computed, pathway-level probabilities were inferred, networks were aggregated, and signaling centrality was calculated on pathway-level networks. Control and disease CellChat objects were then merged within each disease to compare disease-associated changes in interaction number, interaction strength, pathway-level information flow, sender-receiver structure, and ligand-receptor contributions. For pathway discovery and chord visualization, we additionally used a sensitivity-oriented CellChat analysis to capture sparse extracellular matrix-associated signaling. In this analysis, communication probabilities were computed using a 10% truncated mean (type = "truncatedMean", trim = 0.10) with population-size weighting enabled (population.size = TRUE), and interactions were filtered with min.cells = 20. This analysis was used to survey potential ligand-receptor pathway repertoires, whereas disease-control comparisons were based on the main conservative triMean, population.size = FALSE workflow. To evaluate whether major CellChat pathway-level findings were robust to differences in cell numbers across diseases, a balanced stability analysis was performed. CellChat was rerun after downsampling matched condition and cell-type groups across AD, FTD-GRN, and HD. The stability analysis retained T-peri, M-peri, arterial, capillary, and venous endothelial cells, used population.size = FALSE, triMean, and a stricter min.cells = 30, and was repeated across eight bootstrap iterations. Pathway-level disease-control differences were summarized across iterations to assess whether major signaling changes persisted after cell-number balancing.

### Data visualization and statistics

Plots were generated in R using ggplot2, patchwork, ComplexHeatmap, pheatmap, and CellChat visualization utilities. UMAPs were plotted with fixed coordinate ranges within disease datasets where relevant to preserve visual comparability between control and disease conditions. Proportion plots show percentages within each condition. Differential-expression heatmaps display selected significant genes, and pathway-enrichment heatmaps display either negative log10 adjusted P values, normalized enrichment scores, or CellChat-derived pathway-level measures, depending on the analysis layer. For pathway visualizations, displayed terms included top-ranked significant pathways as well as selected biologically relevant pathways of interest, including vascular, extracellular matrix, inflammatory, metabolic, and cell-stress-related pathways. All displayed pathway terms were derived from the corresponding statistical output tables, and pathway selection for visualization was used to aid biological interpretation rather than as an additional statistical test.

Unless otherwise stated, adjusted P values were calculated using the multiple-testing correction implemented in the corresponding tool. Statistical thresholds for differential expression, pathway enrichment, convergence analyses, and cell-cell communication analyses are described above for each analysis layer.

### Software

Analyses were performed in R v4.5.2 using Seurat v5.4.0, Harmony v1.2.4, scDblFinder v1.24.10, MAST v1.36.0, Monocle3 v1.4.26, CellChat v1.6.1, msigdbr v26.1.0, clusterProfiler v4.18.4, gprofiler2 v0.2.4, fgsea v1.36.2, ComplexHeatmap v2.26.1, pheatmap v1.0.13, ggplot2 v4.0.2, dplyr v1.2.1, tidyr v1.3.2, readr v2.2.0, tibble v3.3.1, purrr v1.2.1, patchwork v1.3.2, Matrix v1.7.5, DoubletFinder, and related dependencies.

## Resource availability

### Materials availability

This study did not generate new, unique reagents

### Data and code availability

All data used in this manuscript are publicly available under NCBI GEO accession numbers GSE163577 (Alzheimer’s disease), GSE163122 (GRN-associated frontotemporal dementia), and GSE173731 (Huntington’s disease), and here https://github.com/rustlab1/neurovascular-signature-explorer. We also built an interactive website to search the datasets: https://rustlab1.github.io/neurovascular-signature-explorer/

## Acknowledgements

This work is supported by the funding from USC Dean’s Pilot Funding Award (000092 to R.R.), Epstein Family Breakthrough Alzheimer’s Research Award (000153 to R.R.), National Institute of Neurological Disorders and Stroke (NINDS) (NS141137 to O.B.), and National Institute of Aging (NIA) (AG075798 to O.B.).

## Author contributions

P.M.F. and R.R. contributed to overall project design. P.M.F. and R.R. performed scRNAseq analysis, P.M.F., O.B. and R.R. wrote and edited the manuscript. All authors read and approved the final manuscript.

