## Supplementary material for "Shared and disease-specific human brain vascular signatures in Alzheimer’s disease, frontotemporal dementia, and Huntington’s disease": Supp Figures

### SUPPLEMENTARY FIGURES

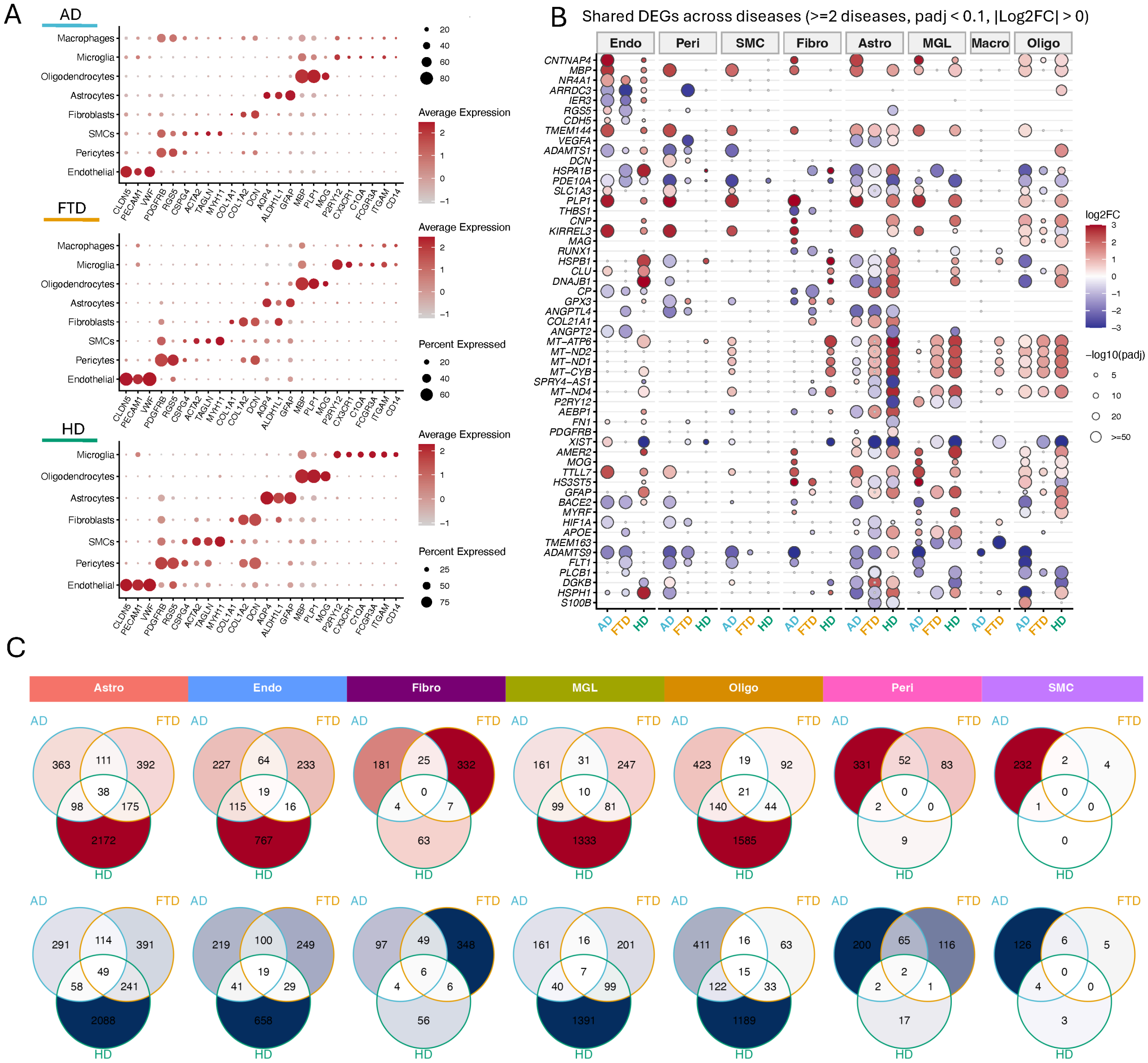


**Supplementary Figure 1. Cell type validation and cross-disease overlap of differentially expressed genes. A,** Dot plots showing expression of canonical marker genes used for cell type annotation in AD, FTD-GRN, and HD datasets. Dot size indicates the percentage of cells expressing each gene, and color intensity represents scaled average expression. **B,** Shared differentially expressed genes (DEGs) across diseases. Genes significant in at least two diseases (adjusted $p < 0.1$, |log2FC| > 0) are shown. Columns represent cell types and rows represent genes. **C,** Venn diagrams showing overlap of significantly upregulated (top) and downregulated (bottom) DEGs (adjusted $p < 0.1$, |log2FC| > 0) across AD, FTD-GRN, and HD within each cell type. Numbers indicate the number of genes in each intersection. AD - Alzheimer's disease, FTD-GRN - frontotemporal dementia caused by GRN mutation HD - Huntington's disease.

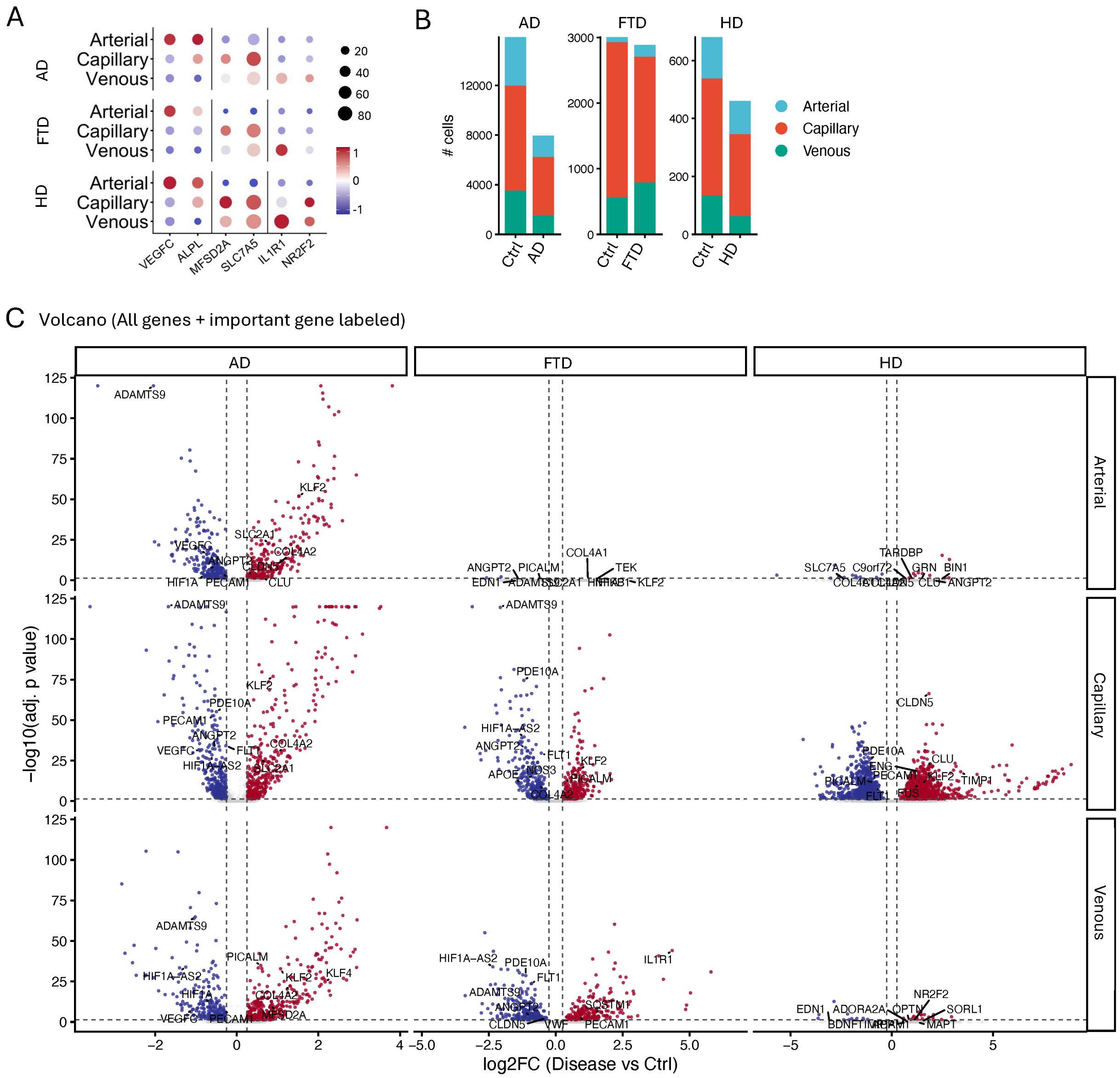


**Supp. Fig. 2: A,** Dot plot of established endothelial zonation markers (VEGFC, ALPL, MFSD2A, SLC7A5, IL1R1, NRP2) across arterial, capillary, and venous segments for AD, FTD, and HD. Dot size represents the percentage of cells expressing each gene; color represents scaled mean expression (red, high; blue, low). Rows are grouped by disease. **B**, Endothelial cell numbers per vascular segment (arterial, capillary, venous) in control versus disease for AD, FTD-GRN, and HD. **C**, Heatmap of segment-specific differentially expressed endothelial genes across diseases, showing scaled disease-control log2 fold changes.

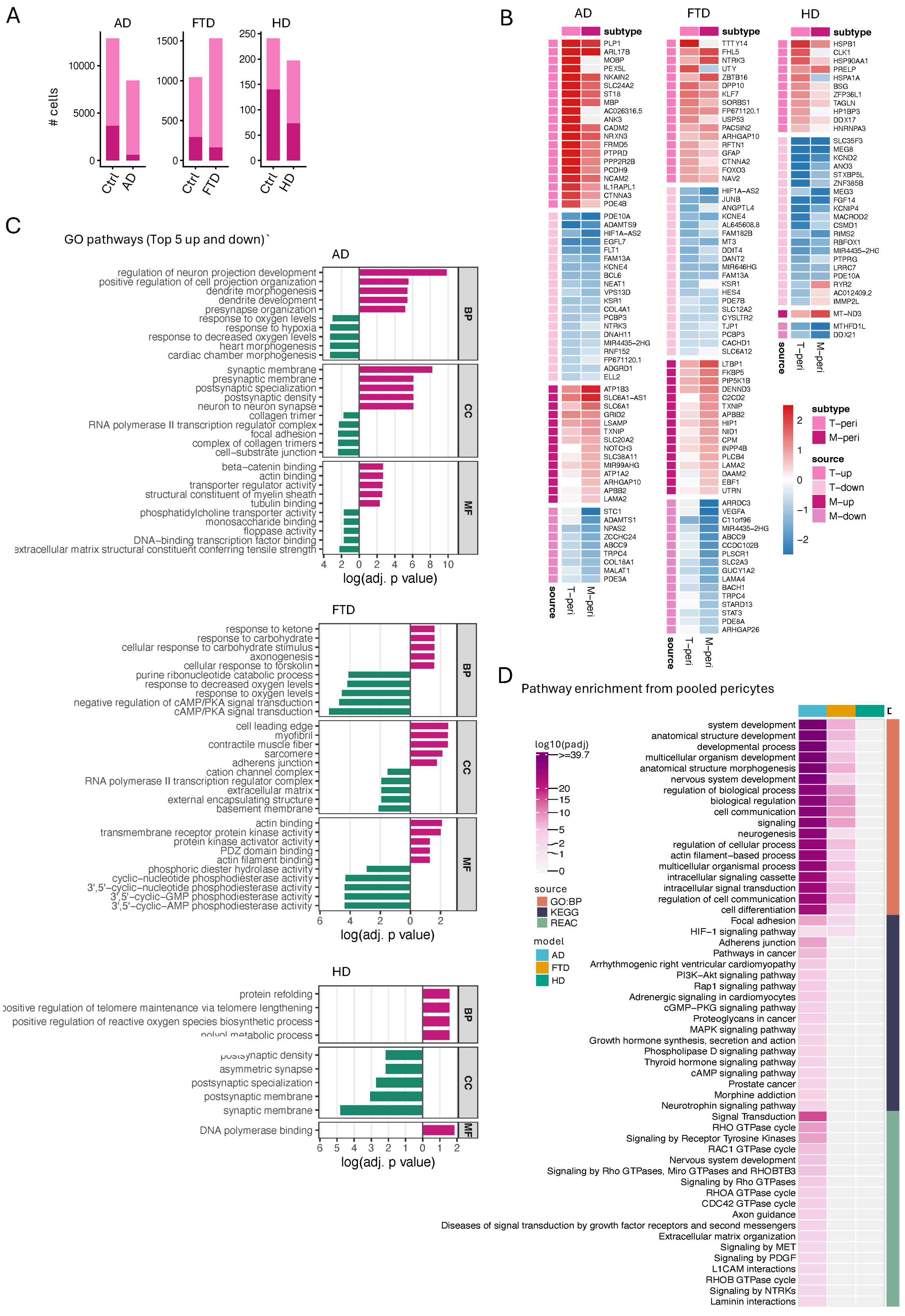

A
AD
FTD
HD
### cells
Ctrl
AD
Ctrl
FTD
Ctrl
HD
B
AD
FTD
HD
subtype
PLP1 ARL17B MOBP PEX5L NKAIN2 SLC24A2 ST18 MBP AC026316.5 ANK3 CADM2 NRXN3 FRMD5 PTPRD PPP2R2B PCDH9 NCAM2 IL1RAPL1 CTNNA3 PDE4B
PDE10A ADAMTS9 HIF1A−AS2 EGFL7 FLT1 FAM13A KCNE4 BCL6 NEAT1 VPS13D KSR1 COL4A1 PCBP3 NTRK3 DNAH11 MIR4435−2HG RNF152 FP671120.1 ADGRD1 ELL2
ATP1B3 SLC6A1−AS1 SLC6A1 GRID2 LSAMP TXNIP SLC20A2 NOTCH3 SLC38A11 MIR99AHG ATP1A2 ARHGAP10 APBB2 LAMA2
STC1 ADAMTS1 NPAS2 ZCCHC24 ABCC9 TRPC4 COL18A1 MALAT1 PDE3A
TTTY14 FHL5 NTRK3 UTY ZBTB16 DPP10 KLF7 SORBS1 FP671120.1 USP53 PACSIN2 ARHGAP10 RFTN1 GFAP CTNNA2 FOXO3 NAV2
HIF1A−AS2 JUNB ANGPTL4 KCNE4 AL645608.8 FAM182B MT3 DDIT4 DANT2 MIR646HG FAM13A KSR1 HES4 PDE7B SLC12A2 CYSLTR2 TJP1 PCBP3 CACHD1 SLC6A12
LTBP1 FKBP5 PIP5K1B DENND3 C2CD2 TXNIP APBB2 HIP1 NID1 CPM INPP4B PLCB4 LAMA2 DAAM2 EBF1 UTRN
ARRDC3 VEGFA C11orf96 MIR4435−2HG ABCC9 CCDC102B PLSCR1 SLC2A3 GUCY1A2 LAMA4 BACH1 TRPC4 STARD13 STAT3 PDE8A ARHGAP26
HSPB1 CLK1 HSP90AA1 PRELP HSPA1A BSG ZFP36L1 TAGLN HP1BP3 DDX17 HNRNPA3
SLC35F3 MEG8 KCND2 ANO3 STXBP5L ZNF385B MEG3 FGF14 KCNIP4 MACROD2 CSMD1 RIMS2 RBFOX1 MIR4435−2HG PTPRG LRRC7 PDE10A RYR2 AC012409.2 IMMP2L
MT−ND3
MTHFD1L DDX21
source
T−peri
M−peri
subtype
T−peri
M−peri
source
T−up
T−down
M−up
M−down
C
GO pathways (Top 5 up and down)`
AD
regulation of neuron projection development
positive regulation of cell projection organization
dendrite morphogenesis
dendrite development
presynapse organization
response to oxygen levels
response to hypoxia
response to decreased oxygen levels
heart morphogenesis
cardiac chamber morphogenesis
synaptic membrane
presynaptic membrane
postsynaptic specialization
postsynaptic density
neuron to neuron synapse
collagen trimer
RNA polymerase II transcription regulator complex
focal adhesion
complex of collagen trimers
cell−substrate junction
beta−catenin binding
actin binding
transporter regulator activity
structural constituent of myelin sheath
tubulin binding
phosphatidylcholine transporter activity
monosaccharide binding
floppase activity
DNA−binding transcription factor binding
extracellular matrix structural constituent conferring tensile strength
BP
CC
MF
log(adj. p value)
FTD
response to ketone
response to carbohydrate
cellular response to carbohydrate stimulus
axonogenesis
cellular response to forskolin
purine ribonucleotide catabolic process
response to decreased oxygen levels
response to oxygen levels
negative regulation of cAMP/PKA signal transduction
cAMP/PKA signal transduction
cell leading edge
myofibril
contractile muscle fiber
sarcomere
adherens junction
cation channel complex
RNA polymerase II transcription regulator complex
extracellular matrix
external encapsulating structure
basement membrane
actin binding
transmembrane receptor protein kinase activity
protein kinase activator activity
PDZ domain binding
actin filament binding
phosphoric diester hydrolase activity
cyclic−nucleotide phosphodiesterase activity
3',5'−cyclic−nucleotide phosphodiesterase activity
3',5'−cyclic−GMP phosphodiesterase activity
3',5'−cyclic−AMP phosphodiesterase activity
log(adj. p value)
HD
protein refolding
positive regulation of telomere maintenance via telomere lengthening
positive regulation of reactive oxygen species biosynthetic process
postsynaptic density
asymmetric synapse
postsynaptic specialization
postsynaptic membrane
synaptic membrane
DNA polymerase binding
log(adj. p value)
D
Pathway enrichment from pooled pericytes
log10(padj)
>=39.7
source
GO:BP
KEGG
REAC
model
AD
FTD
HD
system development
anatomical structure development
developmental process
multicellular organism development
anatomical structure morphogenesis
nervous system development
regulation of biological process
biological regulation
cell communication
signaling
neurogenesis
regulation of cellular process
actin filament−based process
multicellular organismal process
intracellular signaling cassette
intracellular signal transduction
regulation of cell communication
cell differentiation
Focal adhesion
HIF−1 signaling pathway
Adherens junction
Pathways in cancer
Arrhythmogenic right ventricular cardiomyopathy
PI3K−Akt signaling pathway
Rap1 signaling pathway
Adrenergic signaling in cardiomyocytes
cGMP−PKG signaling pathway
Proteoglycans in cancer
MAPK signaling pathway
Growth hormone synthesis, secretion and action
Phospholipase D signaling pathway
Thyroid hormone signaling pathway
cAMP signaling pathway
Prostate cancer
Morphine addiction
Neurotrophin signaling pathway
Signal Transduction
RHO GTPase cycle
Signaling by Receptor Tyrosine Kinases
RAC1 GTPase cycle
Nervous system development
Signaling by Rho GTPases, Miro GTPases and RHOBTB3
Signaling by Rho GTPases
RHOA GTPase cycle
CDC42 GTPase cycle
Axon guidance
Diseases of signal transduction by growth factor receptors and second messengers
Extracellular matrix organization
Signaling by MET
Signaling by PDGF
L1CAM interactions
RHOB GTPase cycle
Signaling by NTRKs
Laminin interactions

**Supplementary Figure 3. Molecular characterization and pathway analysis of pericyte subtypes. A,** Pericyte numbers (# cells) in control versus disease for AD, FTD-GRN, and HD, shown as stacked bars for mesh (M-peri) and tip (T-peri) subtypes. Matrix-associated M-peri decline relative to controls across all three diseases. **B,** Heatmap of canonical pericyte markers and subtype-defining genes across M-peri and T-peri populations in AD, FTD-GRN, and HD. Colors indicate scaled average expression; side bars denote subtype (T-peri, M-peri) and direction of subtype enrichment (T-up, T-down, M-up, M-down). **C,** Gene Ontology (GO) enrichment of disease-associated pericyte DEGs. Top five enriched biological process (BP), cellular component (CC), and molecular function (MF) terms among upregulated (magenta) and downregulated (green) genes for AD, FTD-GRN, and HD. Bar length represents –log10 adjusted p value. **D,** Integrated pathway enrichment of pooled pericyte DEGs across diseases using GO:BP, KEGG, and Reactome. The heatmap displays significantly enriched pathways in AD, FTD-GRN, and HD (columns colored by disease model), with color intensity indicating significance (–log10 adjusted p value). Pathways are grouped by source database (GO:BP, KEGG, REAC) and include cell communication, intracellular signaling, extracellular matrix organization, focal adhesion, HIF-1 signaling, PDGF signaling, and Rho GTPase-mediated pathways. Enrichment was strongest in AD, weaker in FTD-GRN, and minimal in HD.

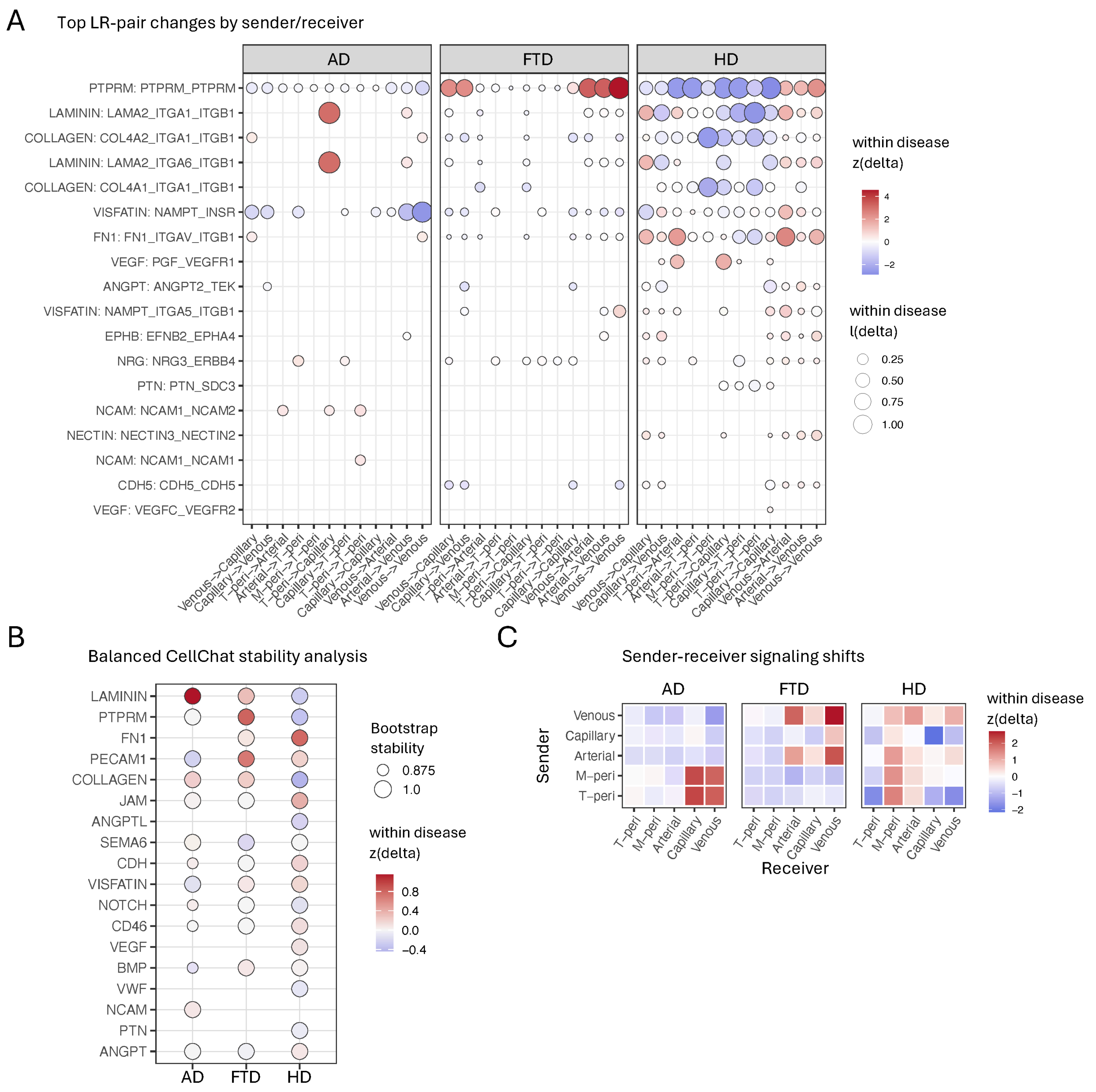


**Supplementary Figure 4. Expanded CellChat ligand–receptor pathway analysis in AD, FTD-GRN, and HD. A**. Ligand–receptor interactions underlying disease-associated signaling changes. Dot plots show selected ligand–receptor interactions contributing to disease-associated CellChat signaling alterations between control and disease conditions across AD, FTD-GRN, and HD. Candidate interactions were selected from the most strongly altered signaling pathways, with the number of ligand–receptor pairs per pathway capped to improve pathway diversity. Sender–receiver cell pairs are shown on the x-axis and ligand–receptor interactions on the y-axis. Dot color indicates the within-disease normalized Disease–Control change in interaction strength, with red denoting increased signaling in disease and blue denoting reduced signaling. Dot size reflects the absolute magnitude of the Disease–Control change. Values were scaled independently within each disease cohort to prevent the larger signaling shifts observed in HD from dominating the visual range. **B**. Stability analysis of disease-associated CellChat pathway changes after cell-number balancing. CellChat was rerun on balanced pericyte–endothelial datasets to assess the robustness of pathway-level Disease–Control differences to unequal cell numbers across cohorts. For each disease comparison, cells were downsampled within each condition and cell type to matched numbers across AD, FTD-GRN, and HD while retaining T-peri, M-peri, and arterial, capillary, and venous endothelial populations. CellChat was performed using the per-cell setting (*population.size*= FALSE), *triMean* averaging, and *min.cells* = 30 across eight bootstrap iterations. Dots represent pathway-level Disease–Control signaling changes. Dot color indicates the median Disease–Control difference across bootstrap iterations, with red indicating increased signaling in disease and blue indicating reduced signaling. Dot size reflects bootstrap stability, defined as the proportion of bootstrap iterations supporting the dominant direction of change. Only the most stable pathway alterations are shown. **C**. Disease-specific rewiring of sender–receiver communication routes.Heatmaps display Disease–Control changes in total CellChat interaction strength between pericyte and endothelial sender–receiver populations for AD, FTD-GRN, and HD. Rows represent sender populations and columns represent receiver populations. Color indicates the within-disease scaled change in total interaction strength, with red representing increased signaling and blue representing reduced signaling in disease relative to control. Values were scaled independently within each disease cohort to highlight disease-specific communication rewiring while minimizing the influence of the globally higher interaction density observed in HD.

### SUPPLEMENTARY TABLES

**Supplementary Table 1. Donor cohort of the three re-analysed human brain snRNA-seq datasets.** Donor-level metadata for all samples included in this study: Alzheimer's disease (8 donors: 4 control, 4 AD; superior frontal cortex), GRN-associated frontotemporal dementia (FTD-GRN; 17 donors: 4 control, 13 FTD-GRN; frontal and temporal cortex), and Huntington's disease (10 donors: 5 control, 5 HD; neostriatum, caudate and putamen). Columns list dataset, donor ID, condition, brain region analysed, sex, age (years), post-mortem interval (PMI, hours), neuropathological stage (e.g., Braak stage and CERAD score for AD), and genotype or additional notes (e.g., APOE genotype). Dashes indicate information not reported in the source study. AD, Alzheimer's disease; FTD-GRN, frontotemporal dementia caused by GRN mutation; HD, Huntington's disease; PMI, post-mortem interval; NR, not reported.
